# Exploring Microbial Dynamics: The Role of Commensals in Healthy and Dandruff-Affected Scalps

**DOI:** 10.1101/2025.03.07.642073

**Authors:** Km. Reeta, Alpana Joshi, Subrata K. Das

## Abstract

The human scalp harbors a diverse microbial community that plays a crucial role in scalp health and disease. This study aimed to analyze the microbiome of both healthy and dandruff-affected scalps to elucidate microbial composition differences and potential associations with dandruff pathology. Samples were collected from three healthy and three dandruff-affected individuals, and the V4 region of the 16S rRNA gene and the ITS1 region were sequenced to characterize bacterial and fungal communities, respectively. Taxonomic analysis identified 5 dominant phyla and 31 genera within the samples, revealing a shift from Actinomycetota dominance in healthy scalps to Basidiomycota dominance in dandruff-affected scalps. Notably, the fungal phylum Ascomycota, including *Malassezia*, was also more abundant in dandruff-affected scalps. This transition emphasizes the intricate interactions among microbial communities and their potential impact on scalp health. These findings suggest specific microbial shifts are associated with dandruff, highlighting the potential for targeted therapeutic strategies that modulate the scalp microbiome to improve scalp health. Further research is needed to elucidate the specific interactions between these microbial groups and their impact on scalp health.

## Introduction

Human skin is a complex microbial ecosystem characterized by intricate interactions among its microbiome and between the host and microbes. The skin provides essential nutrients, such as keratin and lipids, to the colonizing microbes; however, its dry and acidic environment restricts microbial growth. The microbial composition of the skin is influenced by various external factors, including age, gender, hygiene practices, lifestyle choices, immune status, cosmetic use, and medications such as antibiotics and steroids (Smythe & Wilkinson, 2023).

Dandruff is the most common scalp condition, affecting approximately 50% of the population. It arises from factors such as reduced hydration levels, disrupted barrier function, altered cell proliferation, and changes in natural moisturizing factors (Ranganathan & Mukhopadhyay, 2010; Wang et al., 2015). Factors contributing to dandruff include a sedentary lifestyle, oily skin, stress, fatigue, extreme temperatures, and exposure to salty water, infrequent hair washing, and obesity. Dandruff is characterized by hyperproliferation of the scalp epidermis accompanied by itching and redness. It can also be hereditary, known as seborrheic dermatitis (SD). In susceptible individuals, fatty acids penetrate the scalp, leading to inflammation and increased skin cell flaking (Saunte et al., 2020; Naik et al., 2024). Hair loss is a common issue among those with dandruff; in severe cases, it can lead to persistent scalp peeling and may affect other areas, such as the corners of the nose, ears, and eyelids. Microorganisms commonly found in dandruff-affected areas include *Propionibacterium acnes* (*Cutibacterium acnes*), *Staphylococcus epidermidis, Malassezia furfur, Malassezia restricta*, and *Malassezia globosa*. The lipase enzyme produced by *Malassezia* species is considered a critical factor in dandruff development as it catalyzes the production of pro-inflammatory oleic acid, which triggers abnormal peeling of the scalp skin (Grimshaw et al., 2019; Jourdain et al., 2023).

The scalp microbiota consists of a moderately diverse and dense microbial community that inhabits the human scalp. However, the functional role of the scalp microbiome remains largely unknown and warrants further investigation (Saxena et al., 2018). Existing literature provides some insights into how the scalp microbiome influences scalp health and the pathophysiology of related disorders; however, significant gaps remain in our understanding. The identification of bacteria is typically achieved through analysis of the 16S ribosomal RNA (rRNA) gene—a highly conserved genetic marker present in all bacterial species. This gene contains both conserved and variable regions that allow for universal amplification while providing species-specific information. In contrast, fungi and other microeukaryotes are characterized using either the Internal Transcribed Spacer (ITS) region or the 18S rRNA gene. The ITS region exhibits sufficient variability to differentiate between fungal species, while the 18S rRNA gene serves as a eukaryotic counterpart to the bacterial 16S rRNA gene, offering insights into broader eukaryotic microbiota (Yu et al., 2023; López-Aladid et al., 2023; Bartoš et al., 2024).

The application of advanced sequencing techniques, such as amplicon-based surveys and whole-genome shotgun metagenomics, has revolutionized microbiome research, enhancing our understanding of microbial ecology and its implications for human health. However, despite these technological advancements, the mechanisms underlying dandruff pathogenesis remain incompletely elucidated. Factors such as fungal colonization and sebaceous gland activity are implicated in individual susceptibility but require deeper exploration (Tajima et al., 2008; Oh et al., 2014; Tanaka et al., 2016; Park et al., 2017). In the present study, we employed amplicon sequencing to perform a comparative analysis of the microbiota in dandruff-affected and healthy scalp samples. This approach aims to elucidate the potential role of the microbiome in dandruff pathophysiology, offering insights into targeted therapeutic strategies.

## Materials and Methods

### Isolation and Identification of Microflora

Samples were collected from the scalp regions of 6 volunteers, and microorganisms were isolated from 3 non-dandruff scalp (NDS) and 3 dandruff scalp (DS) samples. Scalp surface samples were collected using sterile swabs moistened with sterile saline, ensuring that sterile conditions were maintained throughout the sampling process. A sterile cotton swab soaked in a collection solution (0.15 M NaCl and 0.1% Tween 20) was applied to the scalp surface in a zigzag pattern, covering a total area of 4 cm² without overlapping. The samples were then transferred to four solid media types (Nutrient Agar, MacConkey Agar, Blood Agar, and Luria-Bertani Agar) and four liquid media types (Sabouraud Dextrose Broth, SCD Broth, STO Broth, and Luria-Bertani Broth).

### Genomic DNA isolation and Amplification of Bacterial 16S rRNA and Fungal ITS1 Regions

Genomic DNA was extracted from the samples using the Qiagen PowerSoil kit (Qiagen, MD, USA) following the manufacturer’s instructions with minor modifications. The metagenomic libraries of the fungal and bacterial metagenomics DNA samples were prepared using Illumina Nextera XT sample preparation kit (Illumina Inc., USA). Amplicon libraries were prepared using primers for V4 and ITS1 regions by following the Illumina 16S metagenomics library preparation guide. Amplicon PCR was conducted to amplify the V4 region of the 16S rRNA gene, targeting an approximate length of 270 bp, and to attach the overhang adapter as outlined in the Illumina protocol (2013). The primers used were 533F and 806R (Table 1), which included the overhang adapter sequence along with the specific base sequence for the V4 region. The thermal cycling conditions were set as follows: an initial denaturation at 95°C for 3 minutes, followed by 25 cycles of denaturation at 95°C for 30 seconds, annealing at 55°C for 30 seconds, and extension at 72°C for 30 seconds, concluding with a final extension at 72°C for 5 minutes.

**Table 1.**
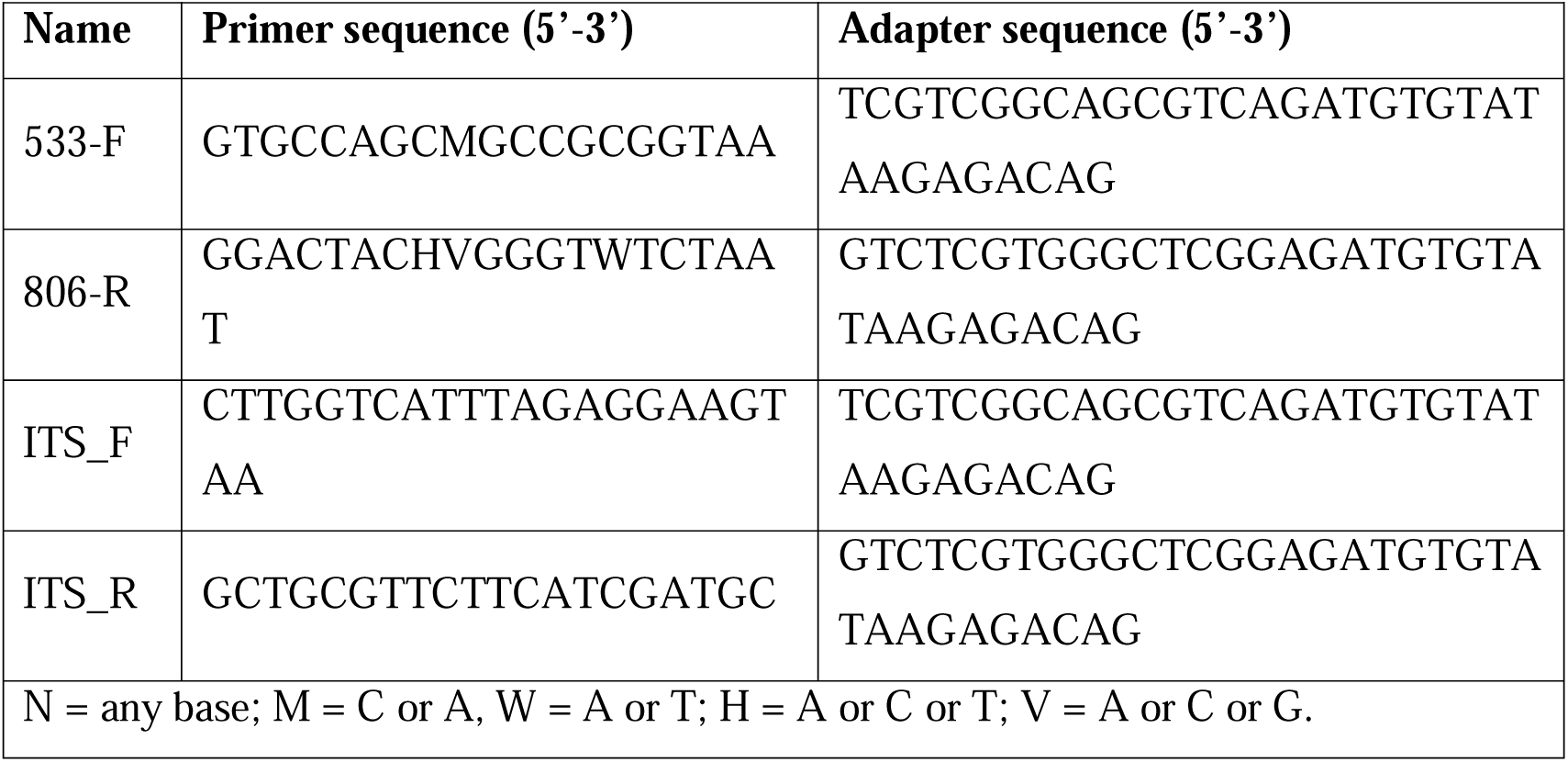
Primers for amplicon PCR.

### Sequencing Data Processing

Base sequences obtained from iSeq 100 sequencing analysis in paired-end format were provided as two FASTQ files for each sample. The raw data were analyzed using the Mothur software (v. 1.48.0), following the MiSeq standard operating procedure (MiSeq SOP) (Kozich et al., 2013).

### Amplification and Sanger Sequencing of ITS and 16S rRNA

The Internal Transcribed Spacer1 (ITS1) and 16S rRNA genes were amplified using standard PCR reactions with universal primer sequences (Table 1). The PCR amplification was performed in a thermal cycler (Applied Biosystems, USA) under the following thermal cycling conditions: an initial denaturation at 94 °C for 4 minutes, followed by 30 cycles of 30 seconds at 94 °C, 40 seconds at 55 °C, and 1 minute at 72 °C. The reaction concluded with a final extension step at 72 °C for 10 minutes. The amplified products were analysed using 1.2% agarose gel electrophoresis, and the PCR products were subsequently purified for sequencing. The cleaned amplicons of ITS and 16S rRNA sequences were bi-directionally sequenced using Sanger’s di-deoxy sequencing method. Sequencing was done following the kit’s protocol (BigDye Terminator v3.1, Applied Biosystems, USA). The same PCR primers have been used for sequencing (Table 1). The reaction was done in ABI 3130xl DNA Analyzers (Applied Biosystems, USA) at Biokart pvt Ltd, Bangalore. Each generated consensus sequence of the forward and reverse sequences was submitted to the Basic Local Alignment Search Tool (BLAST) of the National Centre for Biotechnology Information (NCBI) for a homology search (https://blast.ncbi.nlm.nih.gov/Blast.cgi). Sequences were assembled and edited manually using bioedit v7.05.

### Sequence Alignment and Data Analysis

The sequence alignment was initially performed using the MUSCLE program of MEGA11 with the default alignment parameters for multiple sequence alignment parameters (Tamura et al., 2021). For phylogenetic analysis, we used the Neighbor-Joining tree method with 10,000 bootstraps.

### Statistical Analysis

All experiments were performed in triplicate (n = 3) and results were expressed as mean ± SEM. A one-way analysis of variance (ANOVA) followed by Dunnett’s multiple comparison t-test was used to calculate statistical difference. Statistical significance was considered at *p< 0.05.

## Results and Discussion

### Isolation and Characterization of Microflora from Healthy and Dandruff Scalps

The scalp microbiome was analyzed using three healthy (NDS1, NDS2, NDS3) and three dandruff-affected (DS1, DS2, DS3) samples. Fungal and bacterial diversity was assessed via amplicon sequencing of the 16S rRNA gene for bacteria and the ITS region for fungi, with data processed using Mothur. Sequences were clustered into operational taxonomic units (OTUs) at a 97% similarity threshold, ensuring species-level classification. Taxonomic analysis identified five dominant phyla (four bacterial, one fungal) and 31 genera (28 bacterial, three fungal), highlighting distinct microbial compositions in healthy and dandruff-affected scalps.

### Microbial Community Structure at the Phylum Level in Healthy and Dandruff Scalp Samples

Microbial composition analysis at the phylum level revealed distinct differences between healthy and dandruff-affected scalps (Figure 1). In healthy samples, the predominant bacterial phyla were *Bacillota*, *Actinomycetota*, and *Pseudomonadota*, while *Basidiomycota* represented the dominant fungal phylum. Specifically, in NDS1, the most abundant phyla were *Bacillota* (29.9%), *Actinomycetota* (27.92%), *Pseudomonadota* (24.5%), and *Basidiomycota* (17.75%). A similar trend was observed in NDS2 (*Bacillota* 30.87%, *Actinomycetota* 28.37%, *Pseudomonadota* 24.14%, *Basidiomycota* 16.61%) and NDS3, where *Actinomycetota* was more dominant (38.45%), followed by *Bacillota* (30.52%), *Pseudomonadota* (16.06%), and *Basidiomycota* (14.96%) (Figure 1A). In contrast, dandruff samples exhibited a shift in microbial composition, with a notable increase in *Basidiomycota* abundance. In DS1, the dominant phyla were *Basidiomycota* (54.16%), *Actinomycetota* (32.74%), *Bacillota* (21.39%), and *Pseudomonadota* (15.98%). This pattern persisted in DS2 (*Basidiomycota* 56.84%, *Actinomycetota* 21.42%, *Bacillota* 18.01%, *Pseudomonadota* 15.45%) and DS3 (*Basidiomycota* 48.52%, *Actinomycetota* 25.97%, *Pseudomonadota* 14.43%, *Bacillota* 11.02%) (Figure 1A). The fungal phylum *Ascomycota*, which includes the genus *Malassezia*, was present in both healthy and dandruff samples but was more abundant in dandruff-affected scalps (DS1: 36.24%, DS2: 32.31%, DS3: 38.14%) compared to healthy samples (NDS1: 28.32%, NDS2: 30.86%, NDS3: 31.80%) (Figure 1B).

**Figure 1.**
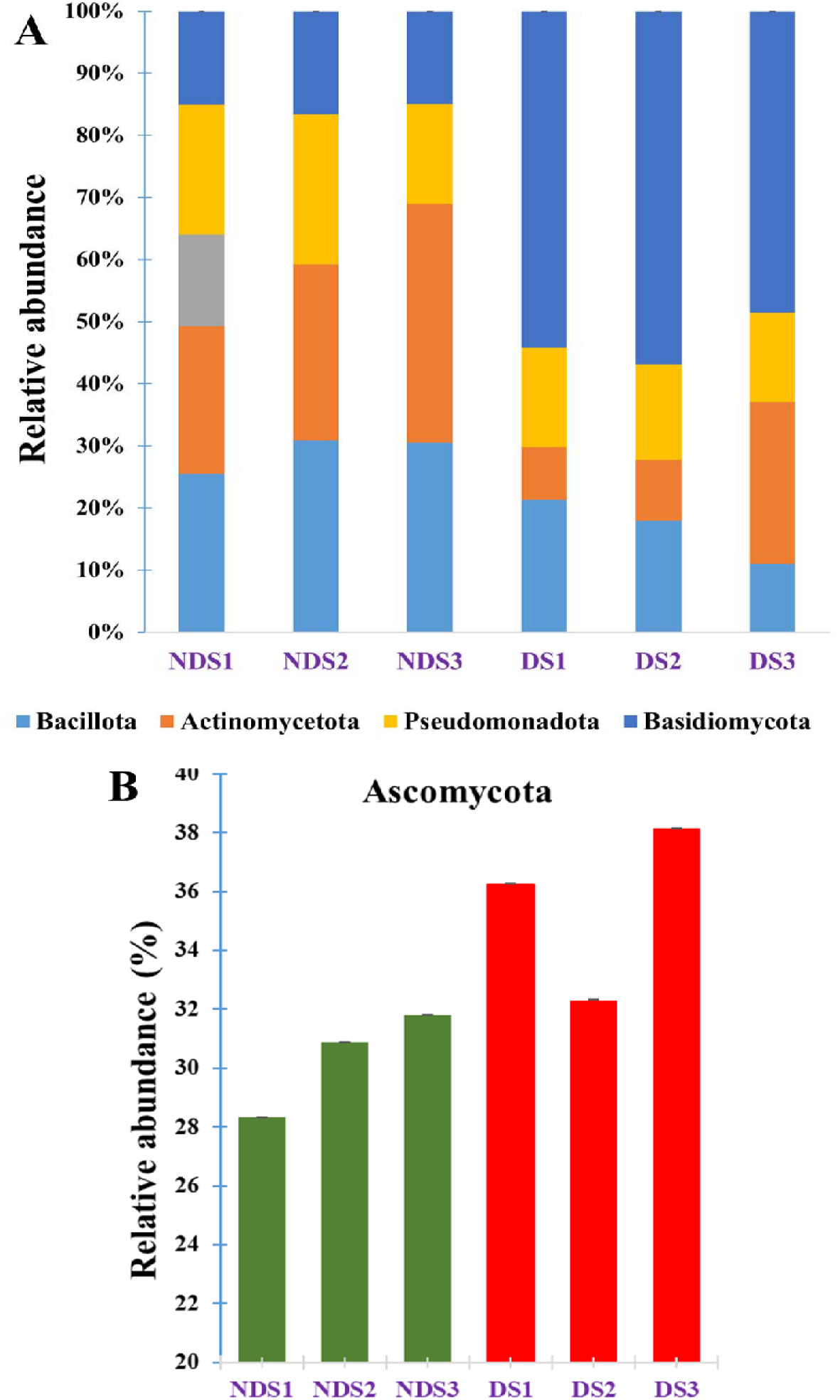
Bacterial (A) and Fungal (B) community structure at the phylum level in healthy and dandruff scalp samples. NDS1 (Healthy sample 1), NDS2 (Healthy scalp 2), NDS3 (Healthy sample 3), DS1 (Dandruff sample 1), DS2 (Dandruff scalp 2), and DS3 (Dandruff sample 3).

Bacteria phylum *Basidiomycota* exhibited significantly higher abundance in dandruff samples (48.52%–56.84%) than in healthy samples, suggesting a potential role in dandruff pathology. While *Pseudomonadota* was detected in both conditions, its varying abundance implies a possible role in scalp health maintenance. Similarly, *Bacillota* was consistently higher in healthy scalps, indicating a potential protective function. *Actinomycetota* was present in both conditions, but its fluctuating levels suggest complex interactions with other microbial communities. These findings highlight distinct microbial shifts associated with dandruff, emphasizing the need for further research on microbial interactions and their impact on scalp health.

### Microbial Community Structure at the Genus Level in Healthy and Dandruff-Affected Scalp Samples

The microbial composition at the genus level was analysed to assess the relative abundance and diversity of microbial taxa across different scalp conditions. In the healthy scalp sample NDS1, the predominant bacterial genus was *Bacillus carotarum* (9.02%), followed by *Brevibacillus laterosporus* (7.05%), *Bacillus cereus* (6.99%), and *Propionibacterium* sp. (6.21%). Other notable taxa included *Bacillus sphaericus* (4.97%), *Erwinia herbicola* (4.89%), *Candida parapsilosis* (4.84%), and *Candida glabrata* (4.40%). Additional bacterial and fungal genera, such as *Enterobacter cloacae* (4.38%), *Staphylococcus aureus* (4.21%), *Cedecea neteri* (3.96%), *Pseudomonas lemoignei* (2.36%), *Klebsiella pneumoniae* (2.30%), and *Staphylococcus epidermidis* (1.64%), contributed to the overall microbial diversity (Figure 2).

**Figure 2.**
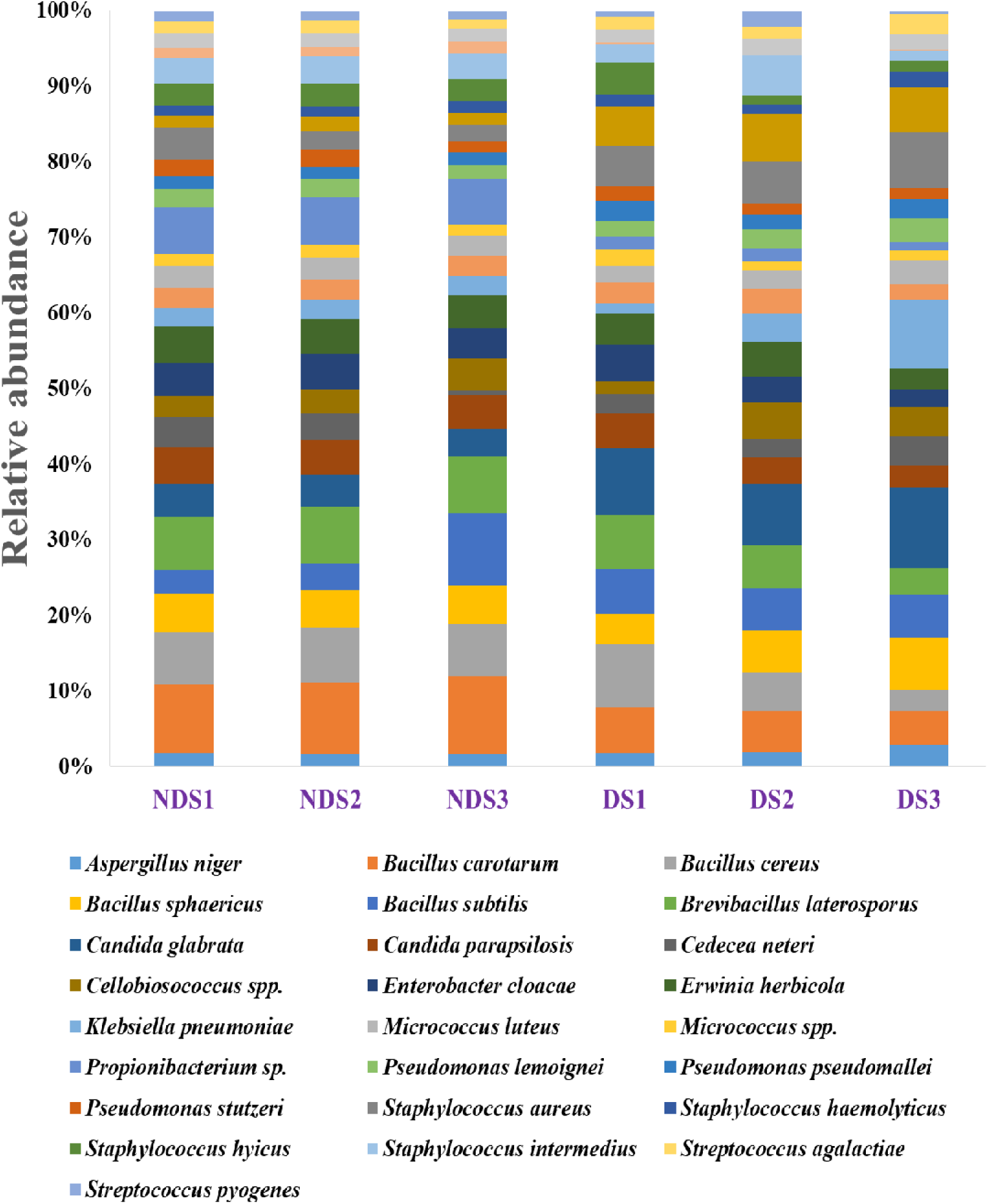
Bacterial community structure at the genus level in healthy and dandruff scalp samples. NDS1 (Healthy sample 1), NDS2 (Healthy scalp 2), NDS3 (Healthy sample 3), DS1 (Dandruff sample 1), DS2 (Dandruff sample 2), and DS3 (Dandruff sample 3).

A similar microbial profile was observed in healthy scalp sample NDS2, where *Bacillus carotarum* (9.41%), *Brevibacillus laterosporus* (7.58%), and *Bacillus cereus* (7.22%) remained dominant. Other prevalent taxa included *Propionibacterium* sp. (6.32%), *Bacillus sphaericus* (4.92%), *Erwinia herbicola* (4.57%), *Candida parapsilosis* (4.60%), *Candida glabrata* (4.16%), and *Enterobacter cloacae* (4.61%). The microbial diversity was further enriched by *Staphylococcus aureus* (2.44%), *Cedecea neteri* (3.53%), *Staphylococcus intermedius* (3.63%), and *Bacillus subtilis* (3.54%) (Figure 2).

In NDS3, *Bacillus carotarum* (10.32%), *Bacillus subtilis* (9.56%), and *Brevibacillus laterosporus* (7.48%) exhibited increased relative abundance. Fungal taxa such as *Candida parapsilosis* (4.53%) and *Candida glabrata* (3.58%) remained consistent with other healthy samples. *Erwinia herbicola* (4.26%) and *Enterobacter cloacae* (4.06%) were prevalent, while *Cellobiosococcus* sp. (4.28%) showed a notable increase. Lower-abundance genera, including *Pseudomonas stutzeri* and *Staphylococcus saprophyticus*, contributed to the microbial diversity (Figure 2). Across all healthy samples, a stable presence of *Bacillus*, *Candida*, and *Propionibacterium* species was observed, with variations in less abundant taxa, highlighting the complex dynamics of the scalp microbiome shaped by bacterial-fungal interactions.

In dandruff-affected scalp samples, a distinct microbial profile emerged. In DS1, the dominant genus was *Candida glabrata* (8.75%), followed by *Bacillus cereus* (8.44%), *Brevibacillus laterosporus* (7.18%), *Bacillus carotarum* (5.94%), and *Bacillus subtilis* (5.91%). Notably, *Staphylococcus aureus* (5.22%), *Staphylococcus epidermidis* (5.15%), *Candida parapsilosis* (4.81%), and *Enterobacter cloacae* (4.80%) were abundant. Additional taxa included *Erwinia herbicola* (4.19%), *Staphylococcus hyicus* (4.09%), and *Bacillus sphaericus* (3.89%), along with lower-abundance genera such as *Pseudomonas pseudomallei* and *Micrococcus* sp. (Figure 2).

Dandruff sample (DS2) exhibited a high prevalence of *Candida glabrata* (8.00%), *Staphylococcus epidermidis* (6.29%), *Bacillus sphaericus* (5.67%), and *Brevibacillus laterosporus* (5.67%). Other dominant genera included *Staphylococcus aureus* (5.62%), *Bacillus subtilis* (5.57%), and *Bacillus carotarum* (5.35%), along with *Staphylococcus intermedius* (5.21%) and *Bacillus cereus* (5.04%) (Figure 2).

Dandruff sample (DS3) displayed an increased abundance of *Candida glabrata* (10.73%), *Klebsiella pneumoniae* (9.12%), *Staphylococcus aureus* (7.28%), and *Bacillus sphaericus* (6.80%). Other prevalent taxa included *Staphylococcus epidermidis* (6.01%), *Bacillus subtilis* (5.78%), and *Bacillus carotarum* (4.52%). Lower-abundance genera such as *Aspergillus niger* and *Streptococcus agalactiae* contributed to microbial diversity (Figure 2).

Fungal diversity analysis revealed that *Malassezia globosa* (26.60%), *Malassezia restricta* (21.79%), and *Malassezia* sp. (18.82%) were the dominant fungal genera in healthy scalp sample NDS1, accompanied by *Aspergillus* sp. (10.75%) and *Cladosporium* sp. (10.48%). Similar trends were observed in NDS2 and NDS3, with slight variations in relative abundance (Figure 3A).

**Figure 3.**
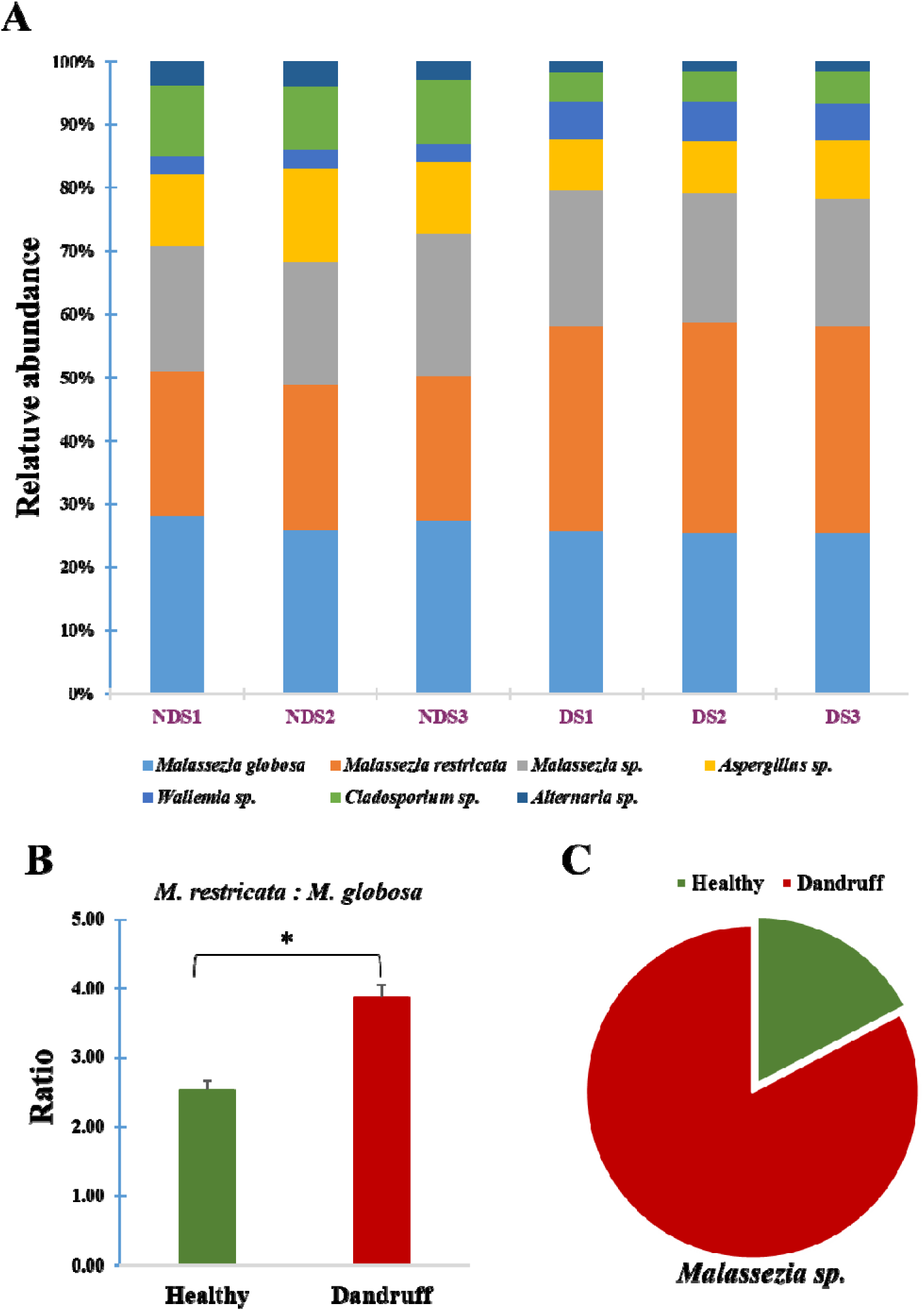
Fungal species diversity in healthy and dandruff scalp samples. (A) Fungal community structure at the genus level in healthy and dandruff scalp samples. (B) Differences in the ratio of *M. restricta* and *M. globosa* in the healthy and dandruff scalp samples (*p ≤ 0.05), and (C) Distribution of *Malessezia sp.* in healthy and dandruff scalp samples. NDS1 (Healthy sample 1), NDS2 (Healthy scalp 2), NDS3 (Healthy sample 3), DS1 (Dandruff sample 1), DS2 (Dandruff scalp 2), and DS3 (Dandruff sample 3).

In dandruff samples, *Malassezia restricta* (32.45%), *Malassezia globosa* (25.64%), and *Malassezia* sp. (21.42%) were significantly enriched in DS1, with a notable increase in *Wallemia* sp. (6.01%) and *Cladosporium* sp. (4.58%). This trend persisted in DS2 and DS3, with *Malassezia restricta* consistently showing higher relative abundance than *Malassezia globosa* (*p* ≤ 0.05) (Figures 3B & 3C).

### Microbial Shifts and Correlation Analysis

A distinct microbial shift was observed in dandruff-affected scalps, characterized by increased *Candida* and *Malassezia* species, alongside higher proportions of *Staphylococcus* and *Klebsiella* species. The elevated *Malassezia restricta* to *Malassezia globosa* ratio in dandruff samples suggests its role in dandruff pathogenesis, emphasizing the need for targeted microbiome modulation strategies.

Correlation analysis revealed a strong positive correlation (*r* = 0.894) between bacterial and fungal populations in healthy scalp samples, indicating a synergistic relationship. However, in dandruff-affected samples, a moderate correlation (*r* = 0.598) suggested microbial imbalance. Specific associations were noted, with *Candida glabrata*, *Staphylococcus epidermidis*, and *Staphylococcus aureus* positively correlating with *Malassezia* species (*M. restricta* and *M. globosa*).

These findings underscore the complex interactions between commensal and pathogenic microbes in maintaining scalp homeostasis. Further research is necessary to elucidate the functional roles of these microbial communities, paving the way for microbiome-based therapeutic interventions to restore scalp health.

### Alpha and Beta Diversity Analysis of Scalp Microbiome in Healthy and Dandruff Samples

Alpha Diversity Analysis: Alpha diversity metrics, including Good’s coverage, observed operational taxonomic units (OTUs), Chao index, and Shannon diversity index, were used to assess the richness and evenness of bacterial and fungal communities in healthy and dandruff-affected scalps. Good’s coverage values exceeded 0.92 for all bacterial samples, indicating high sequencing depth and comprehensive microbial representation. The number of observed OTUs (sobs) ranged from 3297.88 ± 14.29 (NDS2) to 3471.62 ± 34.53 (NDS3) in healthy samples, while dandruff samples exhibited similar values, ranging from 3298.60 ± 18.69 (DS1) to 3450.22 ± 33.08 (DS2). However, species richness, as indicated by the Chao index, was notably higher in dandruff samples (DS1: 11,019.31 ± 524.22, DS2: 12,794.88 ± 393.35, DS3: 11,208.66 ± 323.08) compared to healthy samples (NDS1: 9358.08 ± 405.28, NDS2: 9908.18 ± 367.09, NDS3: 8997.93 ± 290.77), suggesting increased microbial diversity in dandruff-affected scalps. Shannon’s diversity index, which considers both richness and evenness, was higher in dandruff samples (6.04–6.13) compared to healthy samples (5.21–5.42), indicating greater bacterial diversity and a more heterogeneous microbial composition in dandruff scalps (Figure 4A and 4C).

**Figure 4.**
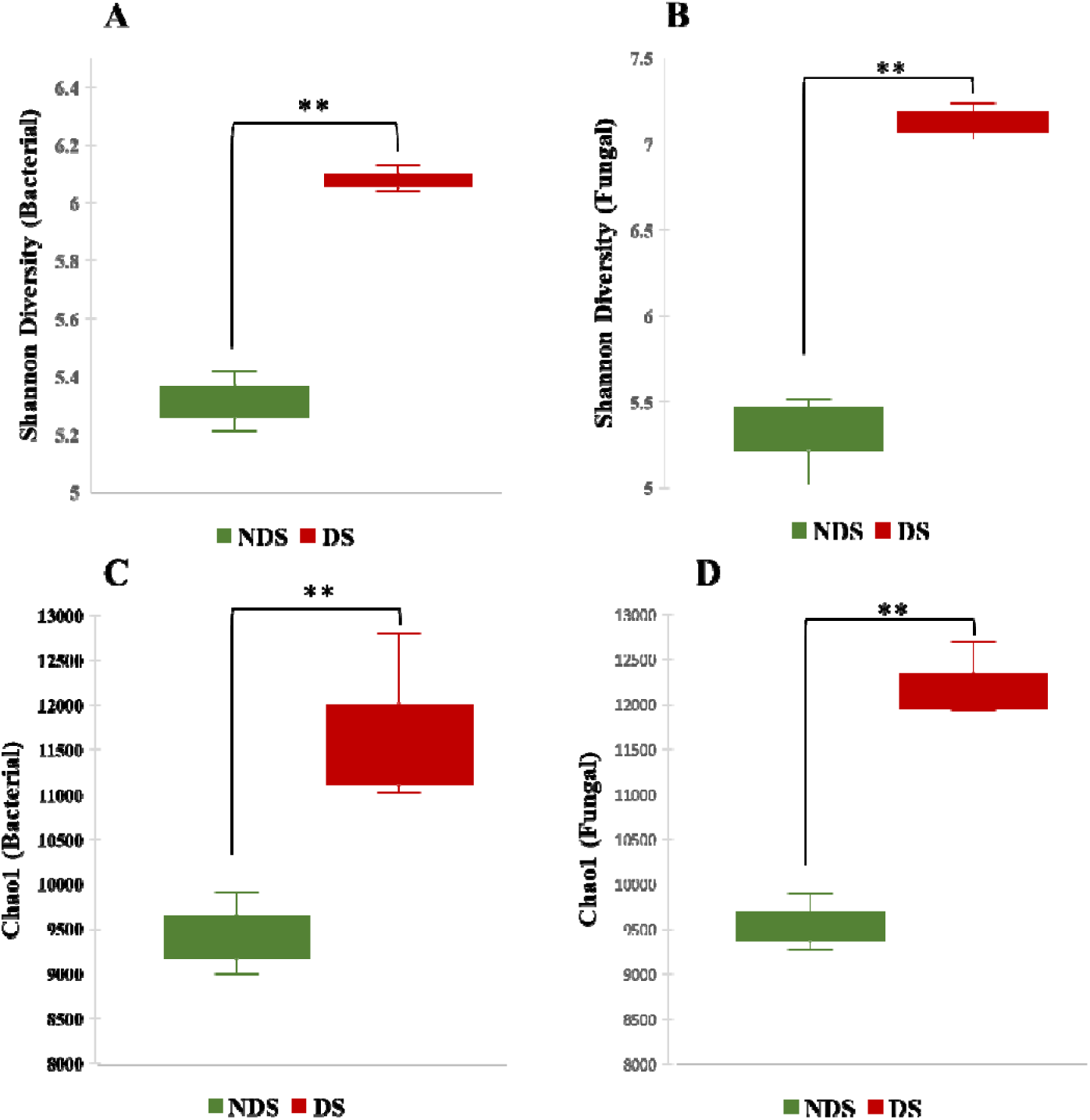
Species diversity of bacterial and fungal microbiome in healthy and dandruff scalp. Shannon diversity index for fungal (A) and bacterial (B) population observed in healthy (NDS: Non-dandruff) and dandruff scalp (**p ≤ 0.001). Shannon diversity index for fungal (A) and bacterial (B) population observed in Non-dandruff (NDS) and dandruff scalp (**p ≤ 0.001).

Similarly, the diversity of fungal communities exhibited a comparable trend. Good’s coverage values were consistently high (>0.94), ensuring reliable sampling representation. The observed OTUs were slightly elevated in dandruff samples (DS1: 4978.60 ± 19.06, DS2: 4870.22 ± 19.68, DS3: 4793.15 ± 25.28) compared to healthy samples (NDS1: 4456.18 ± 15.82, NDS2: 4267.88 ± 18.98, NDS3: 4461.82 ± 24.43).

Moreover, Chao index values were higher in dandruff samples as well, indicating increased fungal species richness. Notably, Shannon’s diversity index was significantly greater in dandruff samples (ranging from 7.03 to 7.24) than in healthy samples (ranging from 5.02 to 5.51), reflecting a more diverse and evenly distributed fungal community on scalps affected by dandruff (Figure 4B and 4D).

The observed increase in bacterial and fungal diversity in dandruff samples suggests a disrupted microbial balance in diseased scalps. Higher Shannon and Chao index values indicate a more complex microbial ecosystem in dandruff conditions, potentially contributing to scalp dysbiosis. The increased presence of certain taxa in dandruff samples, particularly fungal species such as *Malassezia*, and bacterial species associated with pathogenicity, aligns with previous studies linking microbial imbalance to dandruff development. These findings highlight the importance of microbial interactions in maintaining scalp health and suggest that targeting microbial diversity could be a key strategy in dandruff treatment.

Beta-Diversity Analysis: Beta-diversity analysis was conducted to assess variations in microbial community composition across different samples or environments. Common metrics used for studying beta diversity include ThetaYC, Bray-Curtis dissimilarity, and Jaccard (J class) distance. Principal Coordinate Analysis (PCoA) plots revealed distinct clustering patterns for healthy and dandruff samples, indicating high similarity within each group (Figure 5). For bacterial samples, the coordinates on the PCoA plot differed significantly between healthy (e.g., NDS1_B: axis1 −0.298018, axis2 −0.097318) and dandruff samples (e.g., DS1_B: axis1 0.022398, axis2 −0.038269), highlighting unique structural differences in bacterial composition. Similarly, fungal samples exhibited distinct clustering patterns, with healthy samples (e.g., NDS1_F: axis1 −0.621, axis2 −0.0789) differing from dandruff samples (e.g., DS1_F: axis1 0.0476, axis2 −0.187). The Jaccard distance analysis further revealed differences in presence/absence-based metrics among bacterial samples from both groups. Bray-Curtis dissimilarity analysis confirmed significant differences between the microbial communities of healthy and dandruff-affected scalps, supported by distinct coordinates for both bacterial and fungal communities. The distinct clustering patterns observed in the PCoA plots indicate a low degree of similarity in microbial species and their relative abundances between healthy and dandruff-affected scalps. This suggests that dandruff is associated with significant changes in the composition of both bacterial and fungal communities on the scalp. The differences in beta-diversity metrics highlight the unique structural differences in microbial composition between healthy and dandruff samples. These findings support the notion that alterations in scalp microbiota may play a role in the development or exacerbation of dandruff, emphasizing the importance of considering microbial community dynamics in understanding scalp health and disease.

**Figure 5.**
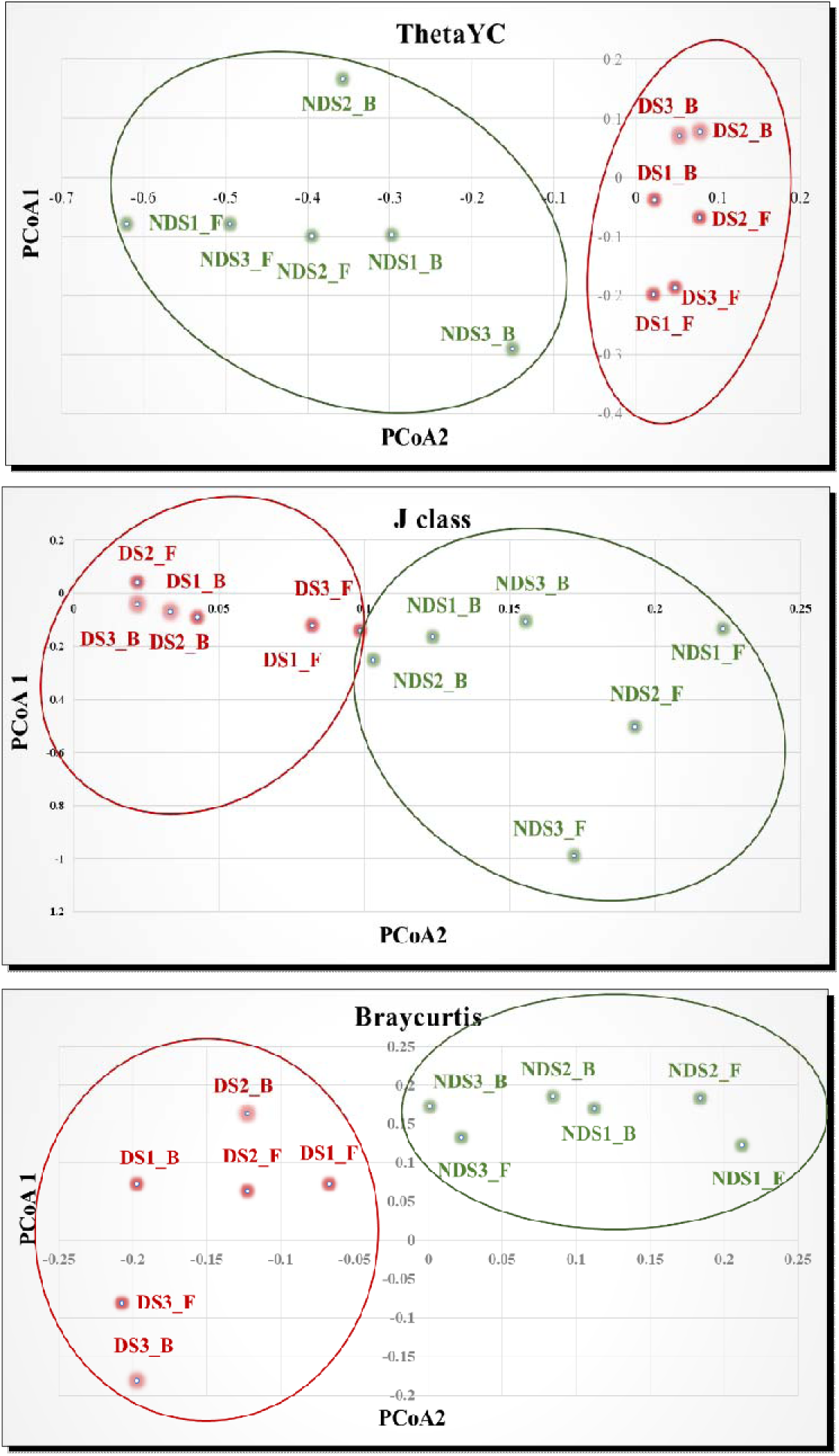
Principal coordinate analysis (PCoA) ordination based on Theta YC, J Class, Bray-Curtis showing significantly different microbial composition among healthy (healthy) and dandruff scalp samples. Bacterial population isolated from healthy (NSD1_B, NSD2_B, and NSD3_B) and dandruff (DS1_B, DS2_B, and DS3_B) scalp samples. Fungal population isolated from healthy (NSD1_F, NSD2_F, and NDS3_F) and dandruff (DS1_F, DS2_F, and DS3_F) scalp samples.

The observed differences in microbial composition could be influenced by various factors, including host intrinsic factors, environmental conditions, and possibly the use of specific treatments or products that affect microbial growth and diversity. Further studies are needed to explore how these changes in microbial communities contribute to scalp health and disease, potentially informing new therapeutic strategies for managing dandruff and other scalp conditions.

### Identification of Specific Microbial Species using 16s rRNA and ITS1 Regions

Genomic DNA was isolated from a randomly selected dandruff sample using a method that leverages the presence of nucleated cells in dandruff, which can serve as a valuable source of genetic material. The extracted DNA was then amplified using specific primers designed to target the bacterial 16S rRNA gene and the fungal ITS1 region. These regions are crucial for identifying microbial species due to their conserved nature and variability among different species. The amplified DNA fragments were subsequently sequenced using Sanger sequencing, a technique known for its high accuracy in generating sequence data. The resulting sequences were compared against established databases using the NCBI BLAST tool to identify microbial species based on their unique genetic signatures. This approach allows for precise identification of bacteria and fungi present in the sample by matching the sequences to known genetic profiles. The results are displayed in Table 2, which outlines the top ten identified species, ordered by their BLASTn scores.

**Table 2.**
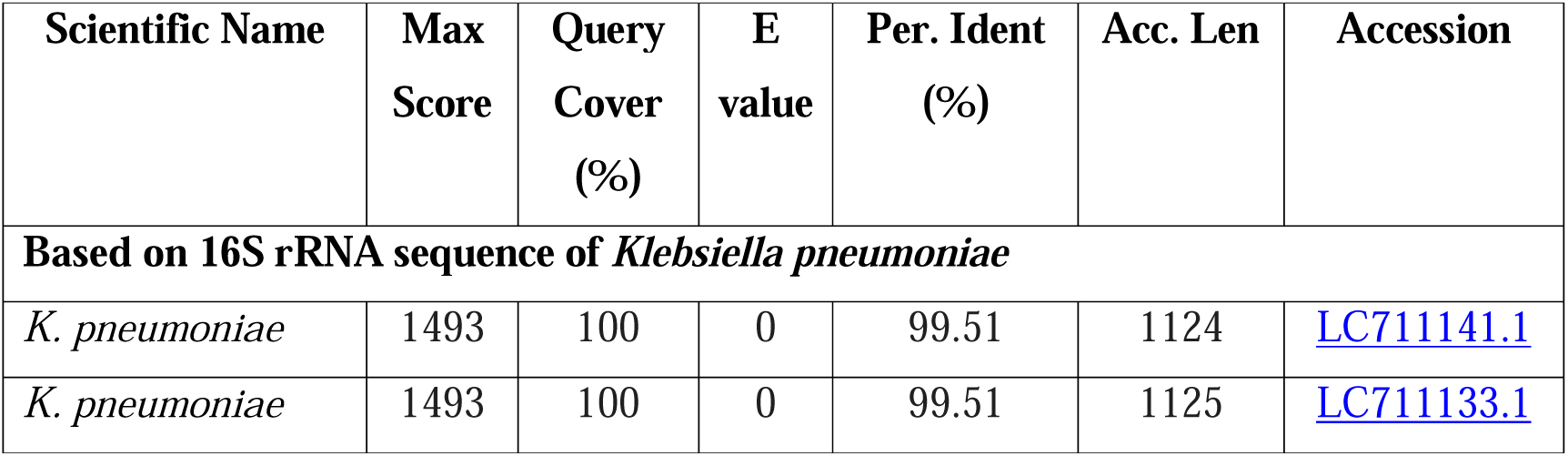
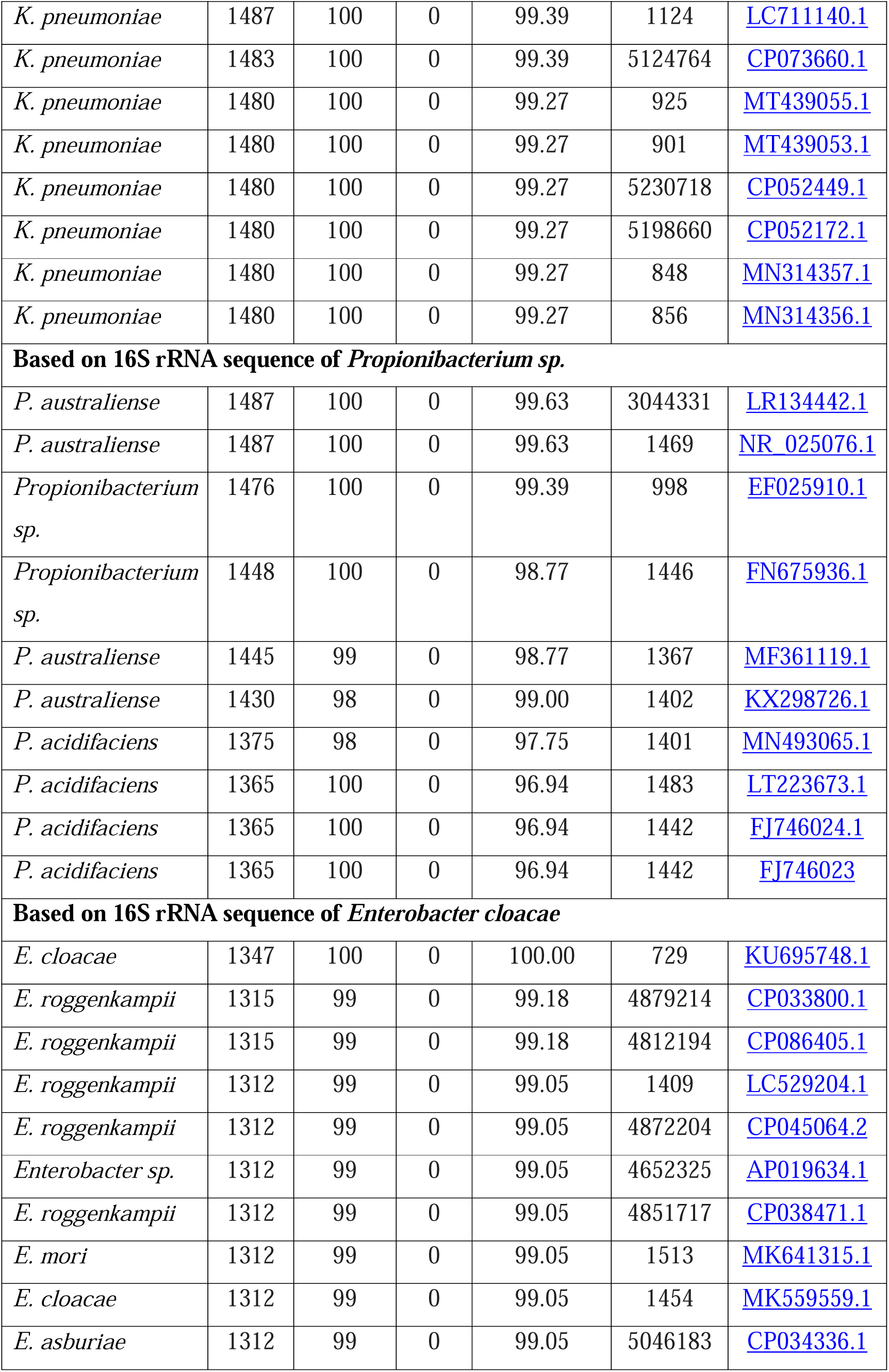
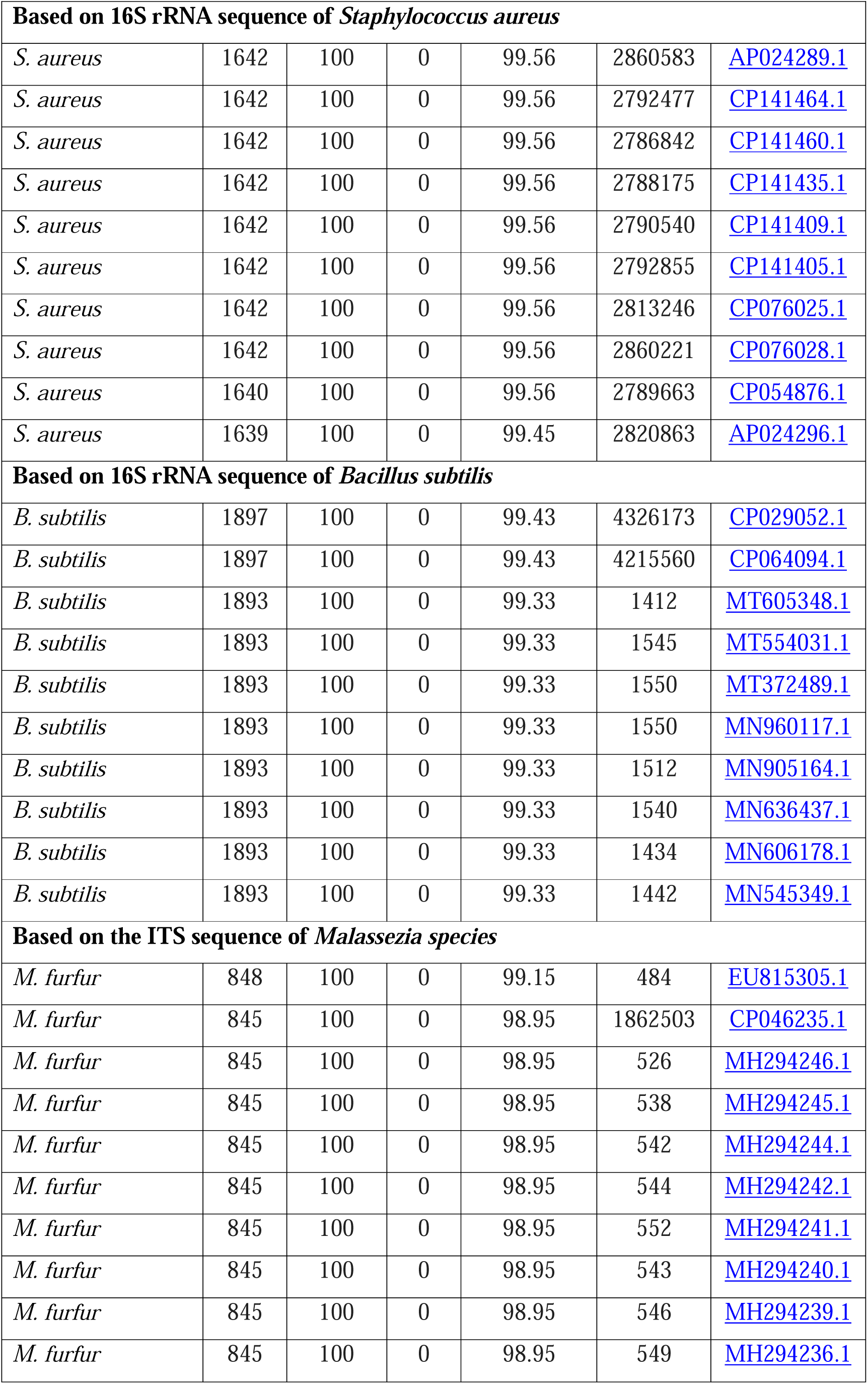
The percentage similarity of sequence based on 16S rRNA and ITS1 regions.

The sequences were aligned using BioEdit software, and the final contigs for each sequence were submitted to the National Center for Biotechnology Information (NCBI) GenBank to obtain accession numbers (OR574403, OR575136, OR574396, OR574399, OR574402, and OR574408). The study identified five bacterial species— *Klebsiella pneumoniae* strain DR-01 (OR574403), *Propionibacterium sp.* strain DR-01 (OR575136), *Enterobacter cloacae* strain DR-01 (OR574396), *Staphylococcus aureus* strain DR-01 (OR574399), and *Bacillus subtilis* strain DR-01 (OR574402) — and one fungal species, *Malassezia furfur* isolate DR-01 (OR574408), in the dandruff-affected scalp sample.

Phylogenetic analysis was conducted using the Maximum Likelihood method combined with the Kimura 2-parameter model to assess the evolutionary relationships among bacterial species based on the 16S rRNA gene. The results showed that *Klebsiella pneumoniae* strain DR-01 (OR574403) exhibited approximately 99% similarity to closely related strains, clustering with 100% bootstrap support. This high degree of similarity indicates a strong evolutionary connection, suggesting that strain DR-01 is closely related to other *Klebsiella pneumoniae* strains within the same clade (Figure 6).

**Figure 6.**
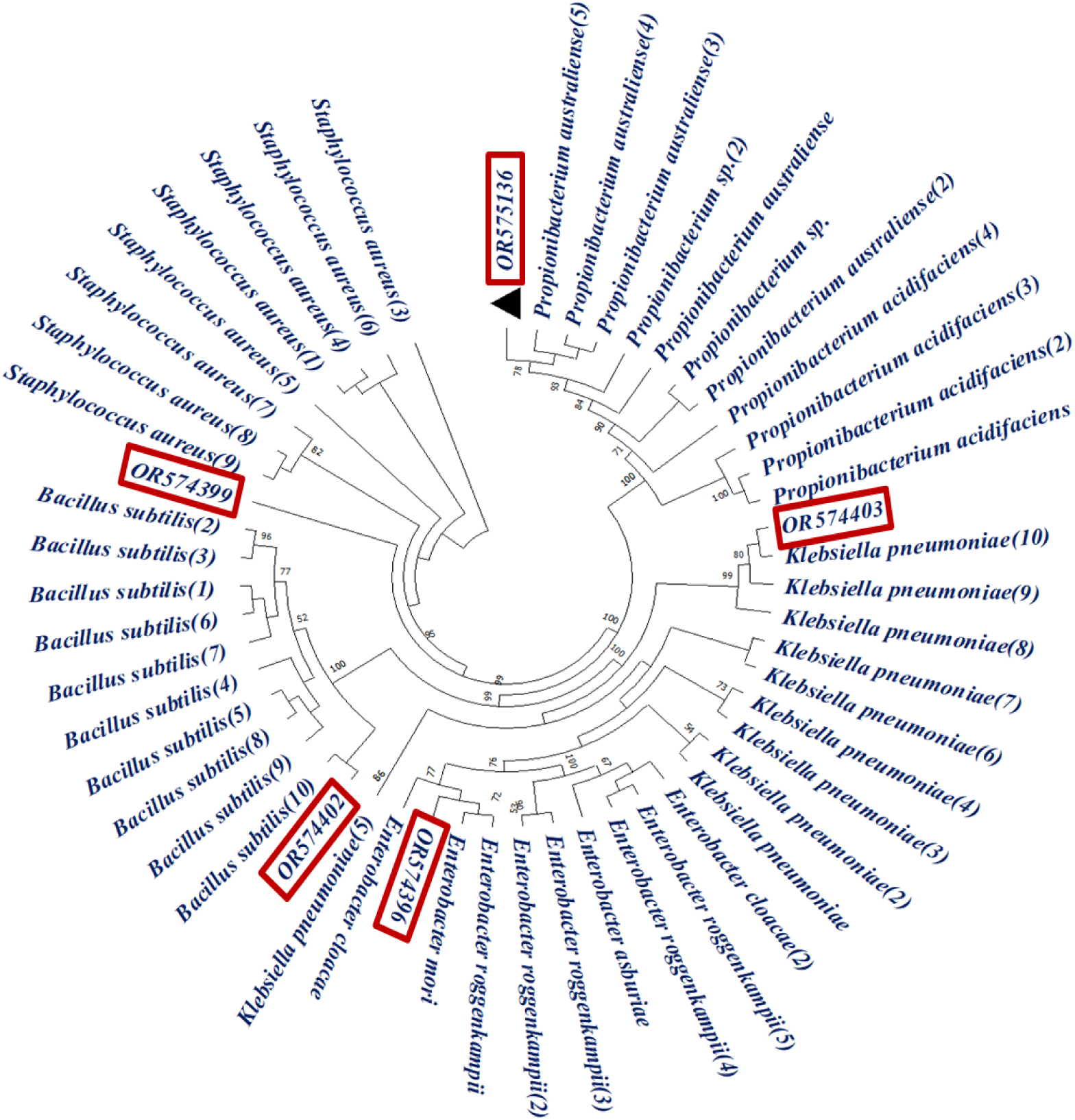
Maximum likelihood phylogenetic tree based on 16S rRNA gene sequences of bacteria. The phylogenetic tree was calculated with MEGA V11 using the maximum likelihood method based on the alignment of the 53 sequences of 16S rRNA gene. The analysis was conducted using MEGA V11, based on the alignment of 53 sequences. Bootstrap values from 1,000 replications are indicated at the branching points, with values <50% was ignored.

Similarly, *Propionibacterium* sp. strain DR-01 (OR575136) showed 96-99% similarity with closely related strains, achieving 100% bootstrap support, indicating a strong evolutionary relationship within its phylogenetic group. *Enterobacter cloacae* strain DR-01 (OR574396) demonstrated 99-100% similarity with related strains and 100% bootstrap support, confirming its close evolutionary ties within the genus. *Staphylococcus aureus* strain DR-01 (OR574399) exhibited approximately 99% similarity to related strains, with 95% bootstrap support, suggesting a robust evolutionary connection. *Bacillus subtilis* strain DR-01 (OR574402) showed approximately 99% similarity to closely related strains, with 100% bootstrap support, highlighting a strong evolutionary relationship within its species. For fungal species, a maximum likelihood phylogenetic tree was constructed using MEGA V11 based on the alignment of eleven ITS gene sequences. The analysis revealed that *Malassezia furfur* strain DR-01 (OR574408) exhibited approximately 99% similarity to closely related strains, emphasizing its close genetic relationship with other *Malassezia* species (Figure 7).

**Figure 7.**
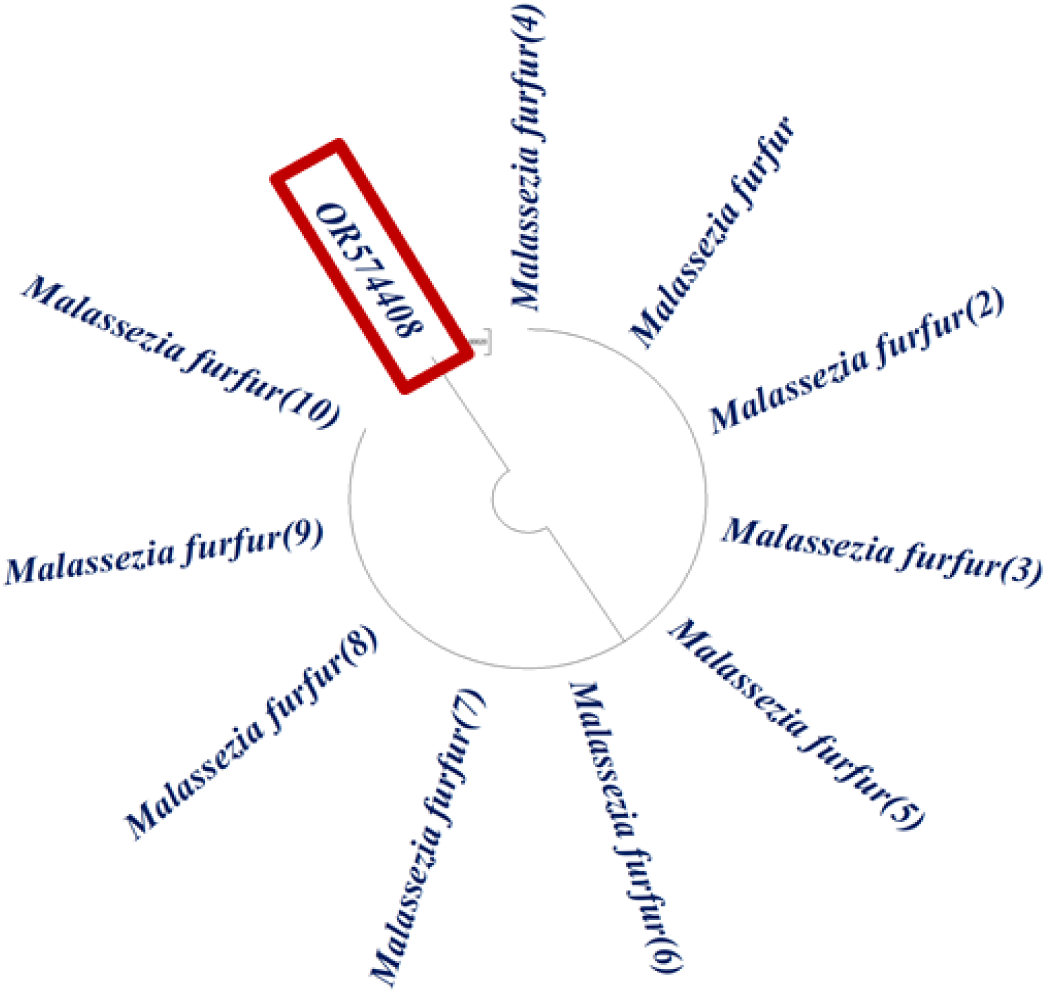
Maximum likelihood phylogenetic tree based on the alignment 11 ITS gene sequences of fungus.

## Conclusions

This study provides valuable insights into the intricate microbial ecosystem of the scalp, highlighting the significant differences between healthy and dandruff-affected scalps. The findings reveal a distinct shift in microbial composition, with a decline in *Actinomycetota* and an increase in *Basidiomycota* and *Ascomycota*, particularly *Malassezia* species, in dandruff conditions. The increased microbial diversity observed in dandruff-affected scalps underscores the complexity of the condition and its potential microbial drivers. The reduction of *Propionibacterium acnes* and the higher prevalence of *Staphylococcus epidermidis* suggest key microbial interactions that may influence scalp health. Furthermore, the observed associations between bacterial and fungal species, such as *Staphylococcus epidermidis* with *Candida glabrata* and *Malassezia restricta*, highlight the need for a deeper understanding of microbial interplay in scalp disorders.

These findings emphasize the potential of microbiome-targeted therapies for dandruff management. Future research should explore probiotic and prebiotic approaches, along with personalized scalp microbiome modulation strategies, to restore microbial balance and improve scalp health. By leveraging microbiome-based interventions, this study paves the way for developing more effective and sustainable solutions for dandruff treatment, ultimately contributing to a better understanding of scalp health dynamics.

